# Baseline Pupil Size Seems Unrelated to Fluid Intelligence, Working Memory Capacity, and Attentional Control

**DOI:** 10.1101/2023.06.22.546078

**Authors:** Veera Ruuskanen, Thomas Hagen, Thomas Espeseth, Sebastiaan Mathôt

## Abstract

Over the past few years, several studies have explored the relationship between resting-state baseline pupil size and cognitive abilities, including fluid intelligence, working memory capacity, and attentional control. However, the results have been inconsistent. Here we present the findings from two experiments designed to replicate and expand previous research, with the aim of clarifying previous mixed findings. In both experiments, we measured baseline pupil size while participants were not engaged in any tasks, and assessed fluid intelligence using a matrix task. In one experiment we also measured working memory capacity (letter-number-sequencing task) and attentional control (attentional-capture task). We controlled for several personal and demographic variables known to influence pupil size, such as age and nicotine consumption. Our analyses revealed no relationship between resting-state pupil size (average or variability) and any of the measured constructs, neither before nor after controlling for confounding variables. Taken together, our results suggest that any relationship between resting-state pupil size and cognitive abilities is likely to be weak or non-existent.

## Baseline Pupil Size Seems Unrelated to Fluid Intelligence, Working Memory Capacity, and Attentional Control

Changes in pupil size are not only brought about by changes in luminance, but also by cognitive activity. For instance, viewing arousing images (Hess & Polt, 1960), memorizing more items during working memory tasks (Kahneman & Beatty, 1966), and performing increasingly difficult mental arithmetic (Hess & Polt, 1964) are all associated with pupil dilation. These so-called psychosensory pupil responses are small in size, but well-established, and are reliable markers of mental effort or arousal (for recent reviews, see Laeng et al., 2012; Mathôt, 2018).

When measuring pupil size, there is an important distinction between task-evoked and baseline pupil size. Task-evoked changes in pupil size are measured during task performance, in response to a specific event, such as the appearance of a stimulus. Much of the research on pupil size has focused on relating the strength of task-evoked changes in pupil size to cognitive factors, such as the amount of mental effort invested in a task. However, more recently the possible relationship between baseline pupil size (as opposed to task-evoked pupil size) and individual differences in cognitive abilities has attracted considerable attention. Baseline pupil size can refer either to a pre-trial baseline, measured during task performance immediately before each trial; or to a pre-experimental baseline, measured in the absence of a task, during a resting state. Here, we will use baseline pupil size to refer to the latter: a pre-experimental, resting-state baseline.

Cognitive abilities that have been suggested to be related to baseline pupil size include working memory capacity (WMC; Aminihajibashi et al., 2019; Heitz et al., 2008; Tsukahara et al., 2016), fluid intelligence (Tsukahara et al., 2016; Tsukahara & Engle, 2021; van der Meer et al., 2010), and attentional control (Unsworth & Robison, 2017). Working memory refers to a cognitive system involved in maintaining, manipulating, and retrieving information relevant to the task at hand, and its capacity is limited (Baddeley & Hitch, 1974; Cowan, 1988; Miller, 1994; Sperling, 1960). Fluid intelligence refers to reasoning and problem-solving abilities (Cattell, 1963; Fukuda et al., 2010). Attentional control refers to the ability to direct attentional resources to the task at hand, and (here) to prevent attentional capture by irrelevant stimuli.

The notion that there might be a relationship between baseline pupil size and cognitive abilities follows mainly from the association between pupil size and activity of the locus coeruleus (LC), such that large pupils indicate high levels of LC activity (Alnæs et al., 2014; Joshi et al., 2016; Murphy et al., 2011, 2014). The LC is a neuromodulatory nucleus located in the brainstem that has projections to the neocortex, including prefrontal areas, and has been suggested to play a modulating role in various cognitive functions, such as attention and behavioral control (Aston-Jones et al., 1999; Aston-Jones & Cohen, 2005; Chandler et al., 2014). Key support for this role of the LC comes from findings showing that people tend to perform better at moments when their pupils are relatively dilated (Keene et al., 2022), presumably because at these moments the LC is relatively active.

Some authors have also linked individual differences in attentional control and WMC to the LC, such that poor performance on tasks involving attention and working memory is linked to dysregulation of the LC (Unsworth & Robison, 2017). Moreover, because of the link between LC activity and baseline pupil size, it has even been hypothesized that differences in these cognitive abilities may be reflected in baseline pupil size as well; in other words, it has been suggested that you can tell someone’s intelligence or working-memory capacity from the size of their pupils at rest (Tsukahara et al., 2016). However, investigations into the relationship between baseline pupil size and cognitive abilities have produced mixed results (see also Unsworth et al., 2021 for a comprehensive review).

The first empirical indications of a relationship between baseline pupil size and WMC (Heitz et al., 2008) as well as between baseline pupil size and fluid intelligence (van der Meer et al., 2010) emerged as incidental findings in studies where the focus was on measurements obtained during task performance. Since baseline pupil size was not the focus of these studies, possible confounding factors such as age and alcohol consumption (Birren et al., 1950; Tryon, 1975) were not considered. Therefore, although these results were suggestive of a relationship, they cannot be taken as decisive evidence.

Following these incidental findings, several studies have provided further evidence for a relationship between baseline pupil size and cognitive abilities. The first large-scale study was conducted by Tsukahara, Harrison & Engle (2016). In a series of three experiments, they found a significant positive correlation between average pupil size and both WMC and fluid intelligence. WMC was measured with a composite score of the operation span, rotation span, and symmetry span tasks; fluid intelligence was measured with Raven’s Advanced Progressive Matrices (RAPM). They accounted for several possible confounding factors such as mental effort, familiarity with the environment, demographic variables (e.g., age), and substance use. When statistically controlling for fluid intelligence, they found that the relationship between pupil size and WMC was non-significant; conversely, when controlling for WMC, the relationship between pupil size and fluid intelligence persisted. Based on this the authors concluded that it is fluid intelligence, rather than WMC, that is uniquely related to baseline pupil size (Tsukahara et al., 2016). However, despite this initial positive finding, the few studies that have since investigated the link between baseline pupil size and fluid intelligence have generally failed to replicate the relationship (Coors et al., 2022; Robison & Brewer, 2022; Robison & Campbell, 2023).

Most subsequent studies have focused on the relationship between baseline pupil size and WMC. However, a number of them, including Aminihajibashi et al. (2019), Coors et al. (2022), Robison et al. (2022), and Unsworth et al. (2019), have failed to replicate the finding of a correlation between average baseline pupil size and WMC. Additionally, a recent meta-analysis concluded that the correlation between baseline pupil size and WMC is not robust (Unsworth et al., 2021). One exception was the study by Aminihajibashi et al. (2019), which found a positive correlation between *variability* in pupil size (indexed by the coefficient of variation [CoV]) and WMC.

Similarly, the few studies that have measured attentional control have generally failed to find a correlation with resting-state baseline pupil size (Robison et al., 2022; Robison & Brewer, 2022; Unsworth et al., 2019). Some studies have demonstrated a relationship with pre-trial baseline pupil size instead, whereby *variability* in baseline pupil size was related to attentional control (but average baseline pupil size was not; Unsworth & Robison, 2017). Overall however, this relationship has received less attention than the correlation between pupil size and both WMC and fluid intelligence.

In spite of the several failed attempts to replicate relationships between baseline pupil size and individual differences in cognitive abilities, methodological differences between studies make it difficult to draw firm conclusions. Tsukahara & Engle (2021) have investigated how these differences may influence results, in an effort to clarify why in their lab they consistently find relationships while many other labs fail to do so. One crucial issue they identified was that different studies measured pupil size under different luminance conditions; this is relevant because a bright laboratory may result in small pupils for everyone, thus reducing variance in pupil size between individuals. With little variance between individuals it becomes difficult to find relationships between variables. Based on both a reanalysis of their own data and a new experiment where luminance conditions were manipulated, Tsukahara & Engle (2021) showed that low variance in the sample of pupil sizes can indeed result in a weak and non-significant correlation between pupil size and WMC.

In addition, based in part on their first study, Tsukahara & Engle (2021) emphasized the importance of the relationship between baseline pupil size and fluid intelligence over other cognitive abilities, such as WMC and attentional control. As mentioned above, many studies have focused on WMC rather than fluid intelligence; however, it seems that only fluid intelligence is uniquely related to pupil size, and that this direct relationship mediates indirect relationships with both WMC (Tsukahara et al., 2016; Tsukahara & Engle, 2021) and attentional control (Tsukahara & Engle, 2021). More generally, all three constructs are highly correlated with each other, with some evidence even suggesting that attentional control largely accounts for the shared variance between WMC and fluid intelligence (Unsworth et al., 2014). Nevertheless, they are still partly distinct (Kane et al., 2005) and could potentially have unique relationships with pupil size.

Following these considerations, here we present the results of two experiments that investigated the relationship between pupil size on the one hand, and fluid intelligence, WMC and attentional control on the other. To account for the relative paucity of studies on this topic, as pointed out by Tsukahara & Engle (2021), our main analysis focuses on fluid intelligence.

Furthermore, the conditions in which our baseline pupil size measurements were obtained closely follow their recommendations. In addition to measuring cognitive abilities, we assessed several possible confounds that are known to affect pupil size (e.g., age, nicotine, alcohol, and caffeine consumption etc.). Our goal was to determine whether previous results on the relationship between pupil size and cognitive abilities are replicable, to help resolve previously observed inconsistencies.

## Methods

### Participants

The results presented here are based on two separate samples of participants, one recruited at the University of Groningen (The Netherlands) and the other at the University of Oslo (Norway).

### Groningen

The sample collected in Groningen consisted of 104 participants, most of whom were undergraduate students at the University of Groningen. These participants were recruited through a web-based system, and they received partial course credit in exchange for participation. The mean age in the sample was 20.05 years, and there were 72 females and 32 males. The majority of the participants were Dutch (35.29 %) or German (32.35 %), while the rest came from various other, mainly European, countries. Participants indicated informed consent by signing a form before the beginning of the study. The study was approved by the local ethics review board (study approval code: PSY1920-S-0178).

### Oslo

The sample collected in Oslo consisted of 122 participants that were recruited through social media. The mean age was 25.56 years, and there were 81 females and 41 males. The study was approved by the local ethics review board (study approval code: 1439337).

## Materials, Apparatus and Procedure

### Procedure

#### Groningen

The experiment began with information and instructions given in both written and verbal form, followed by participants signing the informed consent form. After this, participants filled in a questionnaire on demographic information and possible confounds. Next, a two-minute recording of baseline pupil size was obtained, followed by participants completing the three cognitive tasks described below. Each task took approximately 10 minutes, and the order of the tasks was counterbalanced across participants. Finally, another pupil-size recording was obtained, followed by debriefing. The whole experiment took approximately 45 minutes to complete (Fig 1).

**Figure 1:**
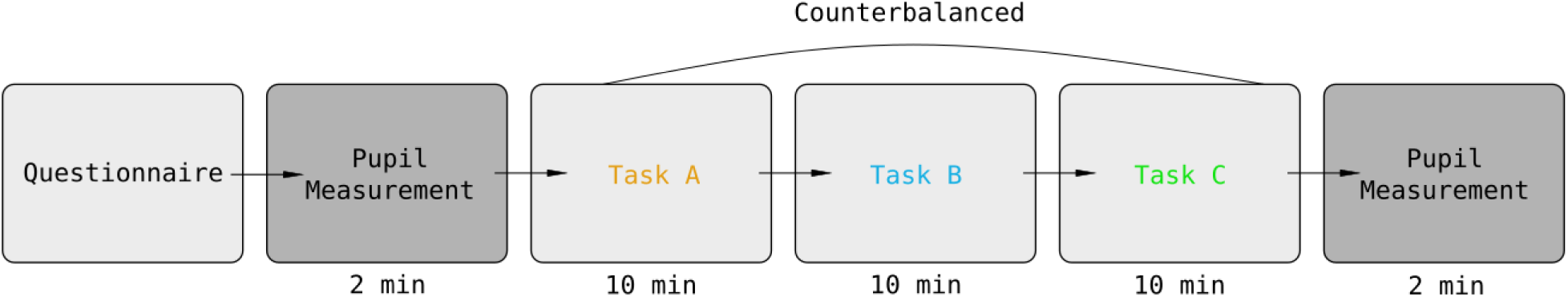
A**n overview of the experimental procedure in Groningen**.

#### Oslo

The experiment began with informed consent and instructions. Participants then filled out a questionnaire on demographic information and possible confounds. Next, a two-minute recording of baseline pupil size was obtained. Participants looked at a fixation cross presented in the middle of a blank screen. Subsequently, participants completed a battery of tasks and questionnaires on cognition and motivation, which will be reported elsewhere. The whole procedure took approximately two hours.

### Pupil measures

#### Groningen

Participants’ pupils were measured twice: at the beginning and end of the experiment. Each measurement was conducted in the absence of a task, and lasted for two minutes. A video-based binocular eye-tracking device (The Eye Tribe, n.d.) was used to record pupil size at a sampling rate of 30 Hz. Pupil size was recorded in arbitrary units. No calibration procedure was applied prior to recording since spatial eye movements were not of interest. A chinrest was used to stabilize head movements and to keep the eye-to-monitor distance constant at 60 cm. All participants were tested in a windowless booth with constant (dim) illumination. During recording participants were asked to fixate on the reflection of their own eyes in the eye tracker. This gives a frontal view of the eye, thus providing the best pupil recording. Further, fixating on the eye tracker provides a dark background for fixation, as recommended by Tsukahara and colleagues (2021) for baseline pupil size measurements. Participants who wore glasses were allowed to keep them on during recording; however, some chose to take them off.

#### Oslo

One measurement was taken, again in the absence of a task, and lasting for 2 minutes. A binocular Remote Eye Tracking Device (R.E.D.; SMI-SensoMotoric Instruments, Teltow, Germany) was used to record pupil size at a sampling rate of 60 Hz. Pupil size was recorded in millimeters of diameter. A chinrest was used to stabilize head movements and to keep the eye-to-monitor distance fixed at 57 cm. The lightning source in the windowless room was in the ceiling, situated slightly behind where the participants were seated, and was set to 50% (rather dim). Participants fixated on a black fixation cross presented on a gray background (RGB: 106, 106, 106).

### Assessment of confounds

#### Groningen

The participants filled out a short questionnaire assessing several demographic variables that may influence pupil size. Specifically, participants reported their age, sex, nationality, handedness, and whether they wore contact lenses (hard or soft) or glasses. In addition, similar to the study by Tsukahara et al. (2016), participants reported: how much sleep they got the night before (in hours); their use of nicotine in the last 10 hours; consumption of alcohol in the last 24 hours; consumption of caffeine in the last 8 hours; and, optionally, use of medication in the past 24 hours. Lastly, participants reported any other factors (if any) they could think of that might influence their pupil size or cognitive abilities, such as memory or attention. Further, the time of the day the experiment took place was recorded.

#### Oslo

Participants verbally reported their age and sex, and whether they had consumed any nicotine or caffeine on the day of the experiment.

### Fluid-intelligence assessment

Both samples of participants completed a matrix task as an assessment of fluid intelligence.

#### Groningen

The task was developed by a research group at the Goethe University in Frankfurt, and is modeled after Raven’s advanced progressive matrices (RAPM; Raven et al., 2000). In this task, participants were shown a three-by-three matrix consisting of eight abstract figures and one empty box (Fig 2a). Underneath the matrix, eight response options were presented. Participants selected the response option that logically belongs in the empty box, thus completing the matrix. The response options were numbered, and participants gave their response by pressing the key corresponding to the number of the chosen option. 10 minutes were given to complete 16 matrices.

**Figure 2:**
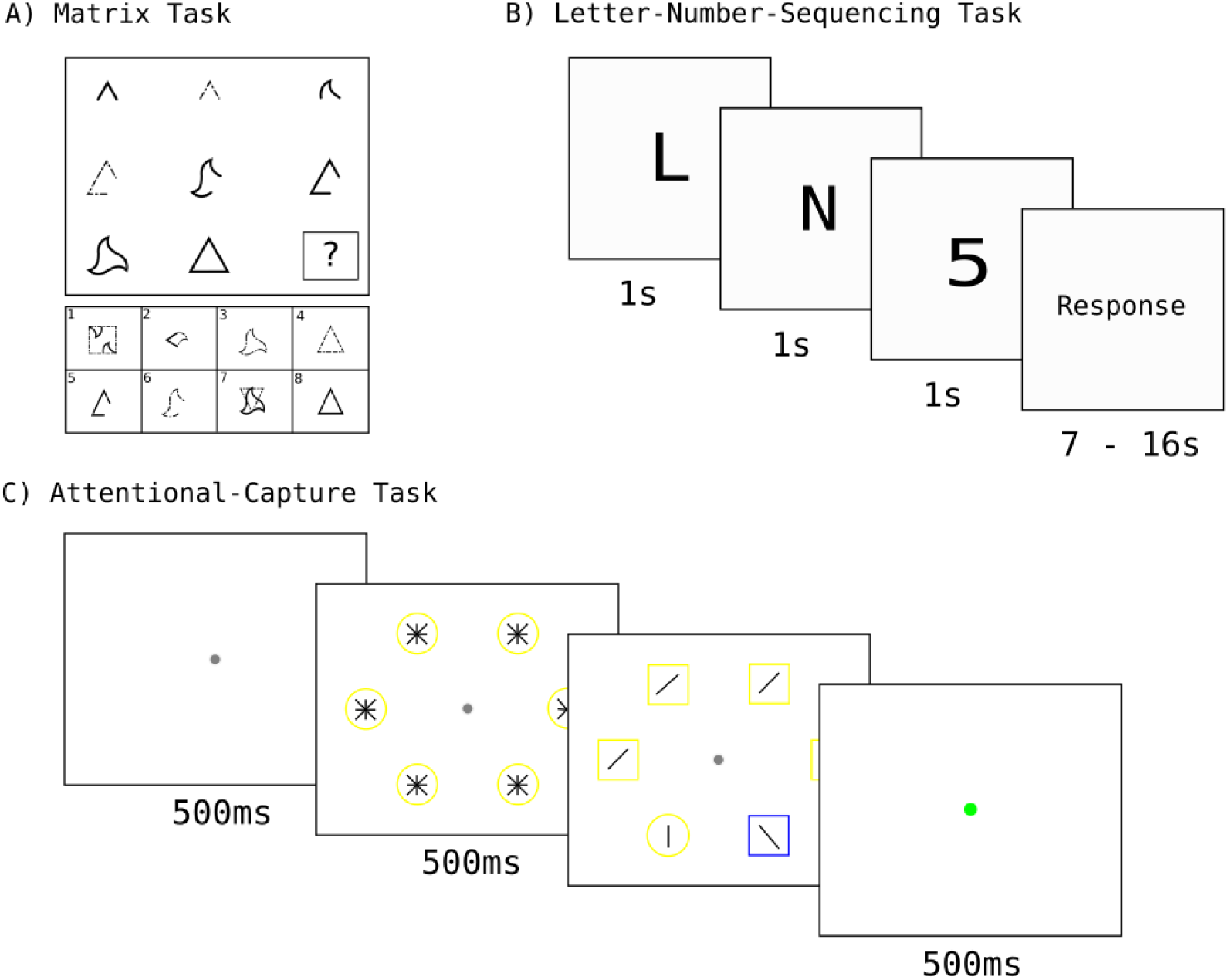
T**h**e **three tasks completed by participants in Groningen**. **A)** An example of a trial of the matrix task used to assess fluid intelligence. **B)** An example of a trial of the Letter-Number-Sequencing task (LNS) used to assess working memory capacity (WMC). **C)** An example of the attentional-capture task used to assess attentional control. *Note*: In the actual task the background was dark and the line segments in the shapes were white. The colors are reversed here for illustration purposes.

#### Oslo

The Hagen Matrices Test (HMT; Heydasch, 2014) was used. The HMT follows the same format as the RAPM, with participants being presented with an incomplete matrix and eight response options. Participants completed eight matrices, and were given maximally two minutes to complete each matrix.

### Other cognitive tasks

#### Groningen

Participants completed assessments of working memory capacity (WMC) and attentional control. All three tasks (including the matrix task described above) were programmed with OpenSesame (version 3.2.8: Kafkaesque Koffka; Mathôt et al., 2012) and presented on a computer screen at a viewing distance of approximately 60 cm. The order of the tasks was counterbalanced such that each possible ordering was completed an equal number of times. The duration of each task was 10 minutes.

WMC was estimated with a letter-number-sequencing (LNS) task (Fig 2b). The LNS is a subset of the Wechsler Adult Intelligence Scale Third Edition (WAIS-III; Wechsler, 1997). In the task, participants see unsorted strings of letters and numbers. They memorize each string and report it with the numbers organized in numerical order, followed by the letters organized in alphabetical order. The length of the strings increases from two to eight. The test is traditionally administered orally (Mielicki et al., 2018). However, to increase ease of administration and analysis, we opted for visual presentation. The procedure and stimuli were the same as in experiment 3-A of a study by Mielicki et al. (2018) that assessed differences in visual and oral administration of the LNS, focusing on language background (bilingual vs. monolingual). They found that visual administration reduces biased results especially for bilingual participants (Mielicki et al., 2018). Given that most participants in the current study speak more than one language, this is ideal. Each character was presented in the center of the screen, in a black monospace font with a font size of 82 px, on a white background, for one second. After the presentation of the entire string, the participants gave their response using the corresponding keys on the keyboard. The time allocated to give a response increased from 7 to 16 seconds (with more time reserved for longer strings), after which the response was considered incorrect. The response times were determined at the author’s discretion. All strings and associated response times can be found in the appendix.

Attentional control was measured with an attentional-capture task that was loosely based on Theeuwes et al. (1992). Participants are presented with six shapes symmetrically organized on a virtual circle on a black background (Fig 2c). On each trial, one of the six shapes is different from the others (a circle among squares or a square among circles). The shapes are blue or yellow in color and contain a line segment. Participants are asked to indicate the orientation of the line in the unique shape by pressing one key on the keyboard for a horizontal line and another for a vertical line. Crucially, on some trials, a distractor shape with a unique color is present, and on others all shapes have the same color. Attentional capture is indicated by the difference in reaction times between distractor-present and distractor-absent trials.

Participants completed 1 block of practice trials, and 8 blocks of experimental trials. A block consisted of 20 trials, making a total of 20 practice trials and 160 experimental trials. Each trial began with a fixation dot presented in the center of the screen for 500ms. Following the fixation, a premask canvas was presented for 500ms. The premask canvas had the target shape and color, with all possible line segments in each shape. Following the premask, the target canvas was presented until a response was given or until three seconds had passed, after which the response was considered incorrect. After each trial, feedback was given in the form of a green (correct) or red (incorrect) dot that was presented for 500 ms. After each block, feedback was given on accuracy and average reaction time during the block.

## Analysis and Results

### Data processing

#### Pupil data

##### Groningen

The preprocessing steps of the pupil data included smoothing with an 11-sample Hanning window and applying a blink-correction algorithm (Mathôt & Vilotijević, 2022). One participant was excluded from the analysis due to a technical measurement error that resulted in an impossible average pupil size value (–1.0). Following preprocessing, the mean and standard deviation over time was determined for each participant separately. Since the Eye Tribe reports pupil size in arbitrary units (as opposed to mm), pupil size was normalized across participants by z-scoring. This was done so that the two samples could be combined for analysis. Further, a coefficient of variation (CoV) of pupil size was computed from the raw data for each participant separately with the following formula: (SD/Mean) * 100. The same formula was used by Aminihajibashi et al. (2019). The CoV of pupil size was also normalized across participants by z-scoring.

##### Oslo

The preprocessing of pupil data was automatically conducted by the SMI software and no additional preprocessing was done. However, the data quality was visually inspected prior to analysis. The SMI R.E.D reports pupil size in mm, but we again normalized the data by z-scoring so that the two samples could be combined. A coefficient of variation (CoV) was computed in the same way as for the Groningen data.

### Task scores

Since the two matrix tasks included a different total number of matrices (Groningen: 16, Oslo: 8), the proportion (as opposed to the number) of correct answers was used for analysis. For the LNS, the number of correct answers was used. No trials were excluded for either of these tasks. For the attentional control task, trials with too short (less than 200ms) or too long (more than 2500ms) reaction times were excluded. This is because in these trials it is likely that the participants were not paying attention to the task. Incorrect trials were also excluded. Next, average reaction times on correct trials were computed separately for distractor-present and distractor-absent trials, and the difference between the two was used as the final score on the task.

### Control variables

The nominal control variables were coded for analysis. For caffeine, nicotine, and alcohol consumption a code of 0 meant the participant had not consumed/used the substance recently and a code of 1 meant they had. Sex was coded such that 1 = female and 0 = male or other.

Handedness was coded such that 1 = right-handed and 0 = left-handed or ambidextrous. Whether the participant wore glasses or contact lenses was coded such that 1 = wears glasses or lenses and 0 = does not wear glasses or lenses. Time of day was rounded to the nearest hour. All data processing was done with scripts written in Python 3 (Rossum & Drake, 2009).

### Analysis

Further analyses were performed using JASP version (0.16.4). One additional participant (the last one) from the sample collected in Groningen was excluded from the analysis to maintain proper counterbalancing across the order of the three tasks. Thus, the final sample size for Groningen was 102 and for Oslo 122. Two sets of analyses were performed. The first analysis focused on the relationship between baseline pupil size (average size and variability) and fluid intelligence, and was conducted using the combined sample of data from both experiments (N = 224). The second analysis investigated the relationships between baseline pupil size (average size and variability) and WMC and attentional control. This analysis was performed on only the sample collected in Groningen, as WMC and attentional control were not assessed in Oslo.

## Distribution of Pupil Sizes

The first step in the analysis was to determine the range and inter-individual variability of recorded pupil sizes. This was done only for the sample collected in Oslo (N = 122), as the pupil measures were in mm, making them comparable with other studies. Average pupil size ranged from 3.01 mm to 7.22 mm with a mean of 4.39 mm and a standard deviation of 0.72 (Figure 3.)

**Figure 3:**
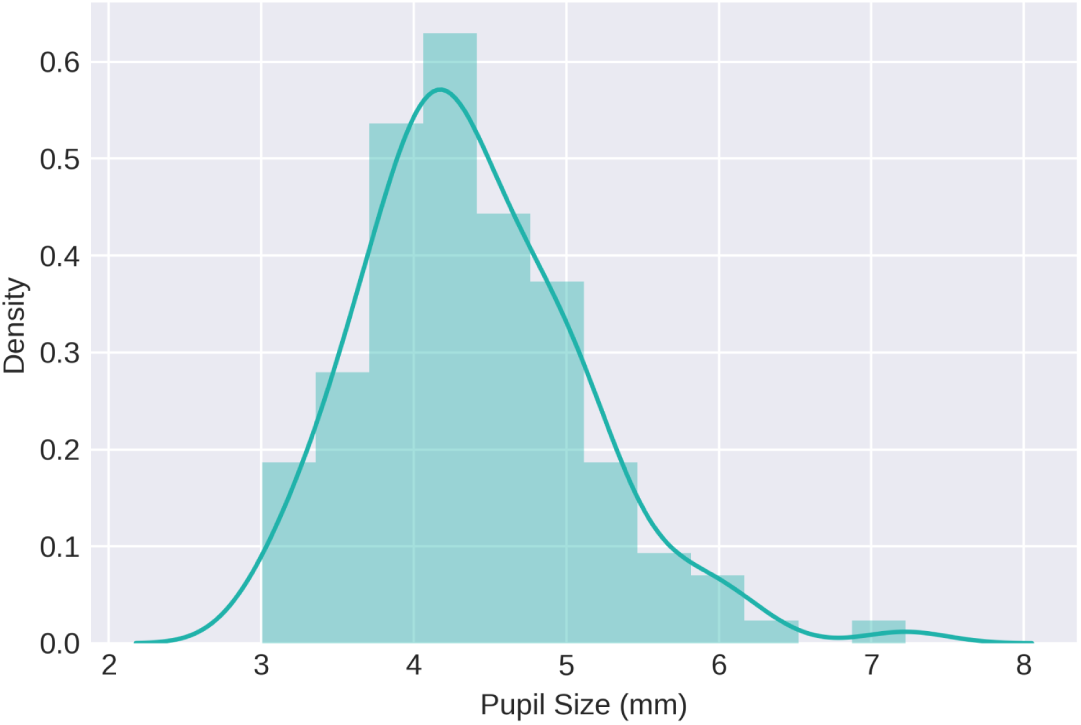
Distribution of resting-state pupil size in mm, as recorded from the sample collected in Oslo (*N* = 122).

The range, mean and standard deviation shown in Figure 3 were extracted from the raw data. However, for the remainder of this section, average pupil size and CoV of pupil size refer to normalized (i.e., z-scored) measures as described above.

## Baseline Pupil Size and Fluid Intelligence

The Pearson correlation between average pupil size and percentages of correct answers on the matrix task was not statistically significant (*r* = –0.01, *p* = 0.83; Fig 4a). A Bayesian Pearson correlation showed support for the null hypothesis (*BF_01_*= 11.6).

**Figure 4:**
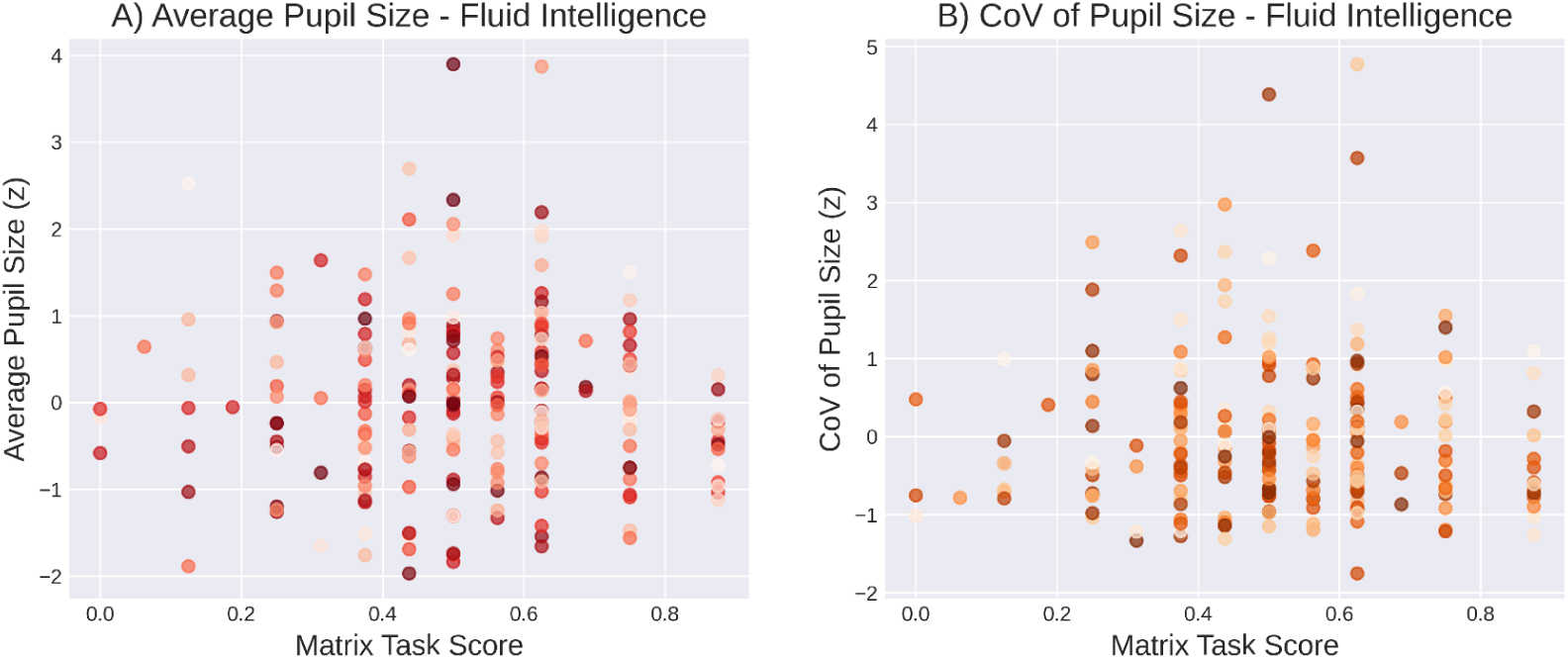
T**h**e **relationship between baseline pupil size and fluid intelligence (as assessed with a matrix task)**. **A)** Average pupil size and fluid intelligence. **B)** Variability in pupil size and fluid intelligence.

Similarly, the Pearson correlation between CoV of pupil size and scores on the matrix task was not significant (*r* = –0.03, *p* = 0.7; Fig 4b). A Bayesian Pearson correlation showed support for the null hypothesis (*BF_01_* = 11.1).

Following the analysis of the correlations between baseline pupil measures and scores on the matrix task, a Bayesian linear regression was performed to investigate the influence of the measured control variables. First, a regression with average pupil size as the dependent variable was conducted. The included covariates were matrix task score, age, sex, caffeine consumption, and nicotine consumption. The coding of the nominal control variables is explained above.

Information on caffeine and nicotine consumption was missing for 11 participants, making the total sample size of this analysis 213. The model that fit the data best included the control variable nicotine but no other covariates (Fig 5). The Bayes factors indicated that the data was 17 times more likely to be observed under this model than the second-best fitting model.

**Figure 5:**
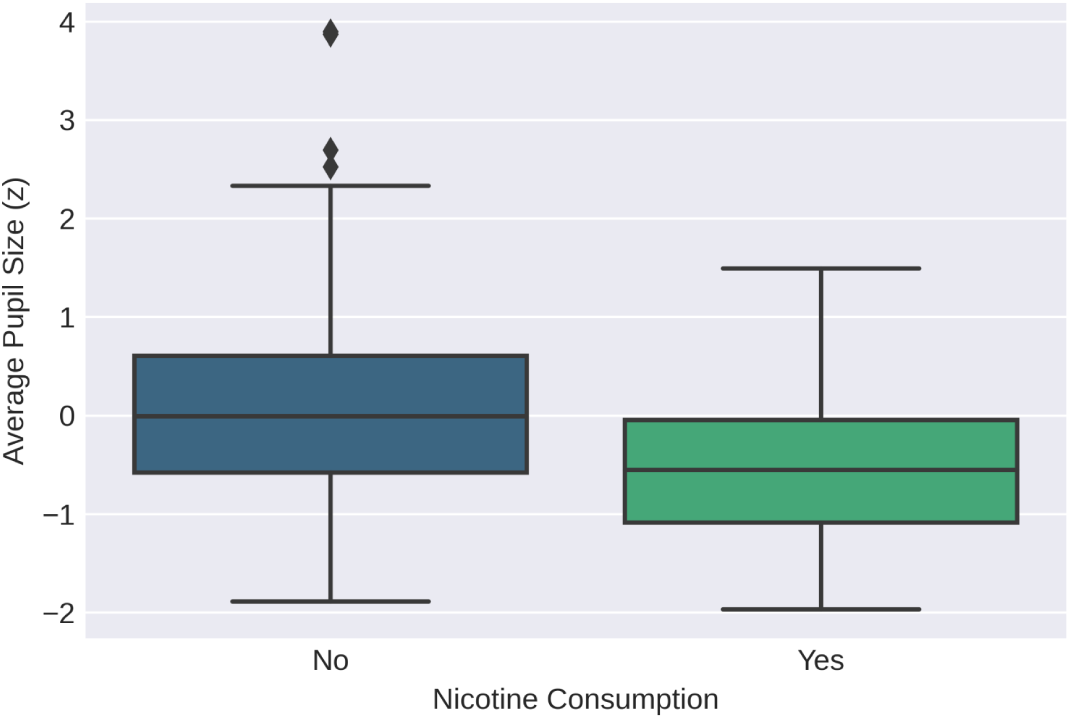
D**i**stribution **of pupil sizes across levels of nicotine consumption in the full sample (0 indicating no nicotine consumption on the day of the experiment and 1 indicating nicotine consumed on the day of the experiment)**.

Next, the same analysis was done with the CoV of pupil size as the dependent variable, and the same covariates as above. The model that fit the data best was the null model.

## Baseline Pupil Size, Working Memory Capacity and Attentional Control

The second analysis focused on the relationships between baseline pupil size and WMC and attentional control. The sample used for this analysis consisted of 102 participants (the Groningen data set).

The correlations between average pupil size and scores on the LNS and attentional control tasks were not statistically significant (*r* = 0.05, *p* = 0.65 and *r* = –0.02, *p* = 0.88, respectively; Fig 6a; Fig 7a). Bayesian correlations showed support for the null hypothesis for both WMC (*BF_01_* = 7.26) and attentional control (*BF_01_* = 8.0).

**Figure 6:**
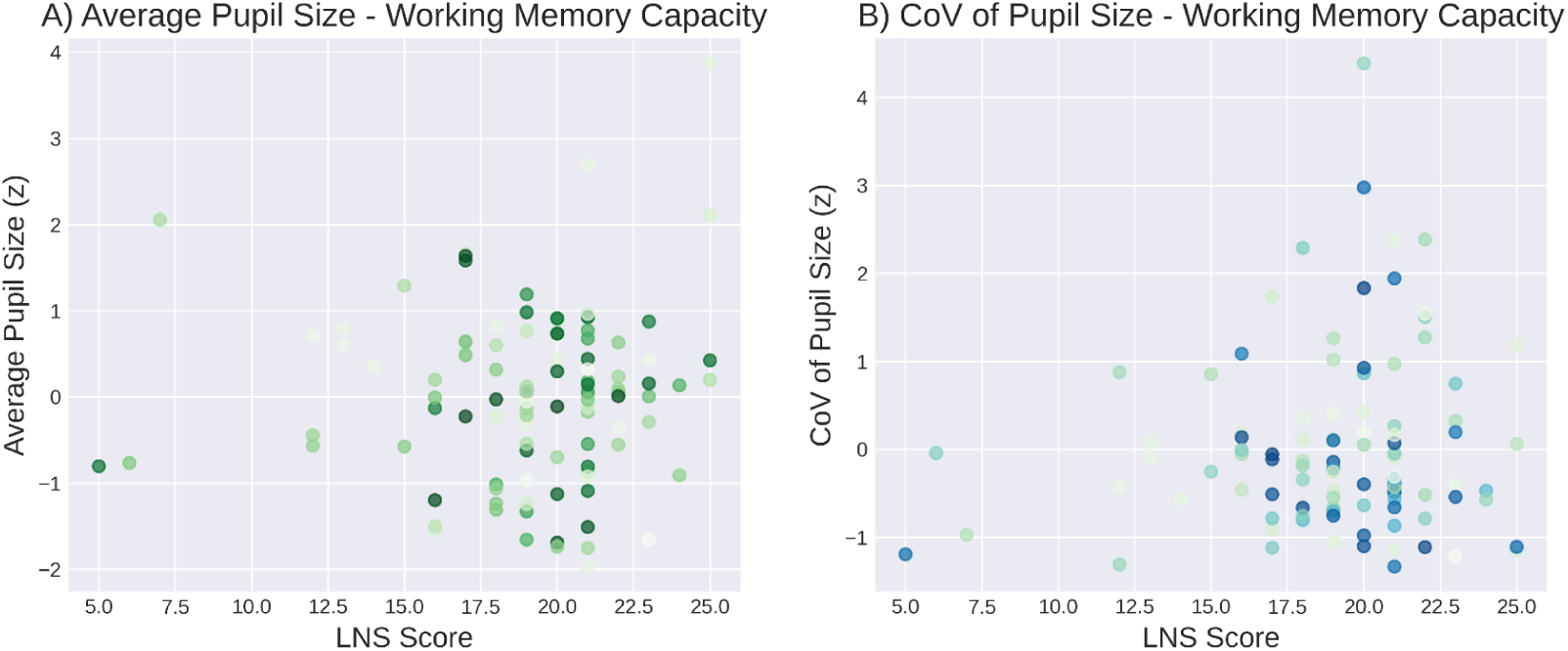
T**h**e **relationship between baseline pupil size and working memory capacity (as assessed with a Letter-Number-Sequencing task)**. **A)** Average pupil size and WMC. **B)** Variability in pupil size and WMC.

**Figure 7:**
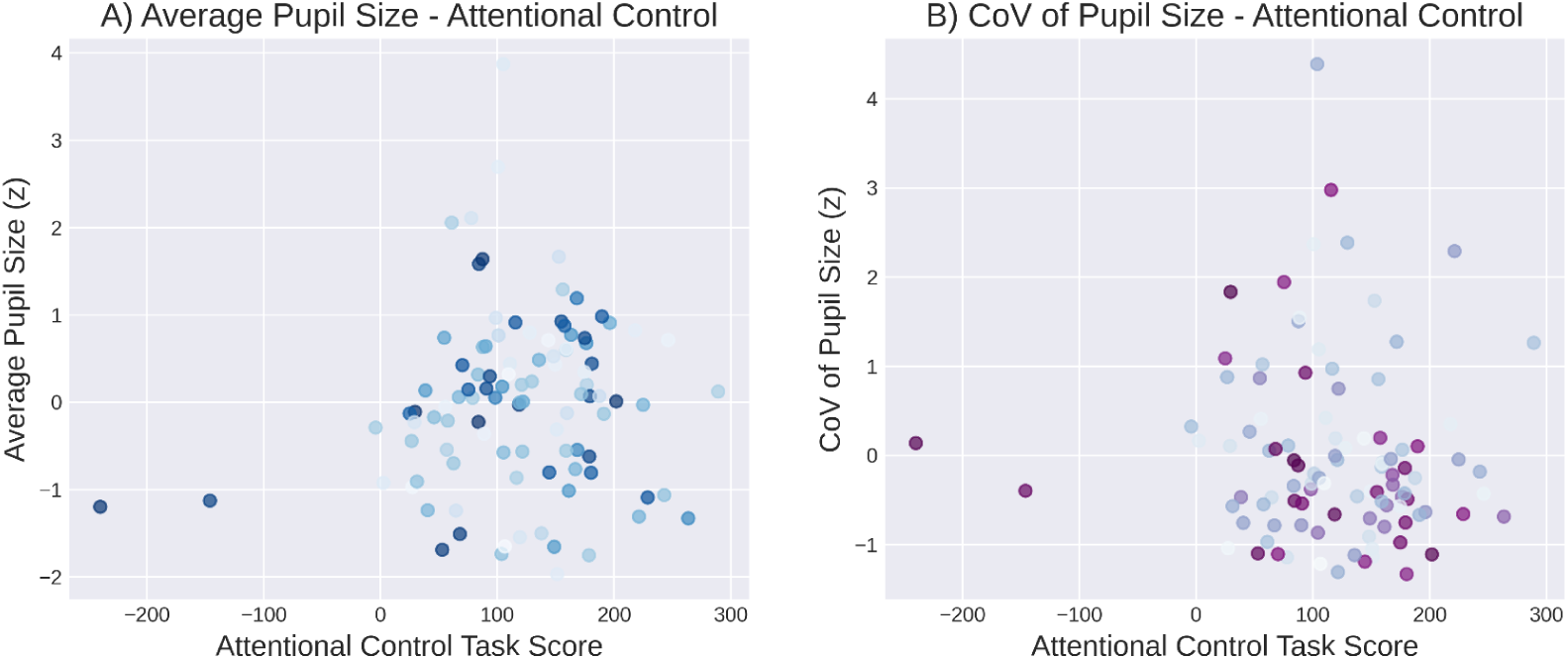
T**h**e **relationship between baseline pupil size and attentional control (as assessed with an attentional-capture task**. **A)** Average pupil size and attentional control. **B)** Variability in pupil size and attentional control.

Similarly, the correlations between the CoV of pupil size and scores on the LNS and attentional control tasks were not significant (*r* = 0.11, *p* = 0.29 and *r* = –0.1, *p* = 0.31, respectively; Fig 6b; Fig 7b). Bayesian correlations showed support for the null hypothesis for both WMC (*BF_01_* = 4.64) and attentional control (*BF_01_* = 4.78)

To investigate the influence of the measured control variables a Bayesian linear regression analysis was performed. The included covariates were the same as above (with score on the matrix task replaced by scores on the LNS and attentional control tasks).

With average pupil size as the dependent variable, the model that fit the data best included the control variable nicotine consumption (Fig 8a) but neither of the tasks. This was followed by a model that included both age (Fig 8b) and nicotine consumption. With CoV of pupil size as the dependent variable, the model that fit the data best was the null model.

**Figure 8:**
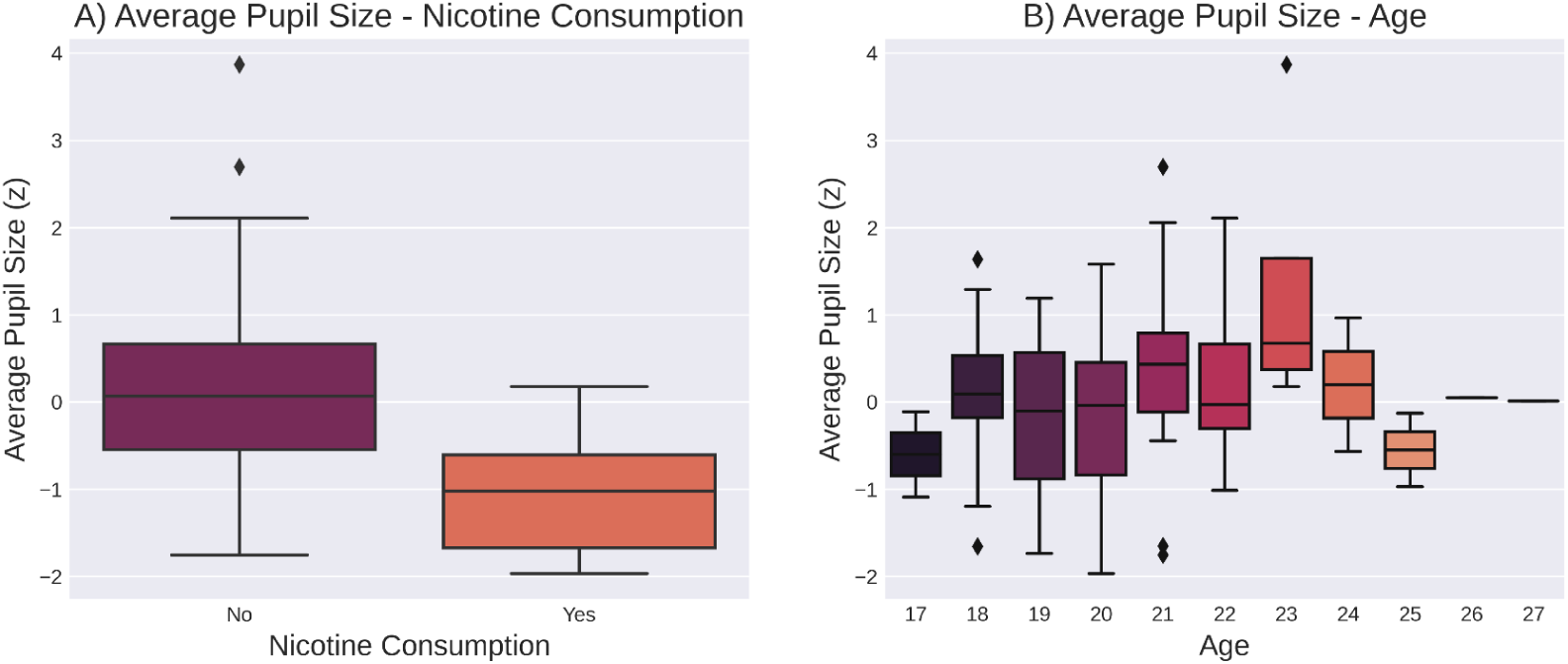
**A**) Distribution of pupil sizes across levels of nicotine consumption in the Groningen sample (0 indicating no nicotine consumption on the day of the experiment and 1 indicating nicotine consumed on the day of the experiment). **B)** Distributions of pupil sizes across age groups.

## Discussion

Here we have presented the results of two experiments that investigated the relationship between resting state pupil size and cognitive abilities, namely fluid intelligence, working memory capacity and attentional control. We did not find any significant relationships between pupil size (average or variability) and any of the measured constructs. In addition, Bayesian analyses provided support for the absence of these relationships in all analyses. We did find that nicotine consumption and age were significantly related to baseline pupil size (see also Wardhani et al., 2020), suggesting that our pupil-size measures were sufficiently reliable to pick up individual differences.

Crucially, we failed to replicate previous findings despite following several of the recommendations given by Tsukahara & Engle (2021) to ensure ideal conditions for observing relationships between baseline pupil size and cognitive abilities. Tsukuhara and Engle (2021) suggested that the relationship between pupil size and fluid intelligence is more robust than that between pupil size and other cognitive abilities, such as WMC (Tsukahara & Engle, 2021; Unsworth et al., 2021). Therefore, and unlike many previous studies, we measured fluid intelligence in addition to WMC and attentional control, and focused our main analysis on fluid intelligence. However, despite taking the suggestions by Tsukuhara and Engle (2021) to heart, we did not manage to replicate previous findings (see also Coors et al., 2022; Robison et al., 2022; Robison & Campbell, 2023).

Another possible problem in previous studies was that pupil size was measured in overly bright settings, leading to constricted pupils and reduced inter-individual variability in pupil size. In both of our experiments, participants fixated on a dark (black or gray) background during the baseline pupil measurement. This ensures a sufficiently large average pupil size, and also sufficient inter-individual variability. Further, participants in Groningen fixated on the eye tracker rather than the screen; this gives a frontal view of the eye, giving the best pupil recording. The mean and standard deviation of pupil sizes recorded in our Oslo sample were 4.39 mm and 0.72, respectively. This falls between the values reported by Tsukahara et al. (2016; mean 5.92 mm, standard deviation 1.09) and Unsworth et al. (2019; mean 3.21, standard deviation 0.49).

In addition to background brightness, a notable strength of our measurement conditions was using a chin rest to stabilize the participant’s head and to keep the eye-to-monitor distance fixed across participants. In contrast, Tsukahara et al. (2016) did not use a chin rest, which may influence recorded pupil sizes. For instance, highly motivated participants might sit closer to the eye tracker, leading to a larger average pupil size and more inter-individual variability.

Considering the recommendations by Tsukahara et al. (2021), one limitation of our study was using only a single task to measure each construct. Different measures may tap into different aspects of the underlying construct, making it advisable to use multiple measures and derive the common variance across them for a more precise measurement (Ackerman & Hambrick, 2020). However, the specific task used is unlikely to influence results if the underlying correlation is robust. In addition, the WMC and fluid-intelligence tasks employed in our experiments were the same or similar to those used in previous studies (Aminihajibashi et al., 2019; Tsukahara et al., 2016), further supporting the conclusion that the observed results are not a consequence of the tasks used here. Moreover, Robison & Campbell (2023) did use multiple tasks to measure fluid intelligence and were still unable to replicate the link between baseline pupil size and fluid intelligence.

Another concern raised about previous studies is their lack of diversity, as most of the samples have consisted mainly of North American college students, with few exceptions (Coors et al., 2022; Robison et al., 2022). In this study, we recruited some participants through social media, resulting in a more ability-diverse subsample (with a mean age of 25.56 years) than the typical undergraduate student sample. Additionally, the majority of our participants came from various European countries, which could introduce differences in latent variables such as motivation or arousal level in testing situations when compared to the typical North American college student sample (for a more in-depth discussion, see Aminihajibashi et al., 2019).

In conclusion, we do not find evidence for a relationship between baseline pupil size (average or variability) and cognitive abilities (fluid intelligence, WMC, and attentional control), despite rigorous measurement conditions and a sufficient (though modest compared to some other studies) sample size. Out of the variables that we measured, only age and nicotine consumption were significantly related to baseline pupil size. Our results suggest that the relationship between baseline pupil size and cognitive abilities is weak and not particularly robust.

## Open practices statement

The data and analysis code are available on the OpenScience Framework (https://osf.io/vf9sb/). We did not pre register either study.

